# Using eDNA to biomonitor the fish community in a tropical oligotrophic lake

**DOI:** 10.1101/375089

**Authors:** Martha Valdez-Moreno, Natalia V. Ivanova, Manuel Elías-Gutiérrez, Stephanie L. Pedersen, Kyrylo Bessonov, Paul D.N. Hebert

## Abstract

Environmental DNA (eDNA) is an effective approach for detecting vertebrates and plants, especially in aquatic ecosystems, but prior studies have largely examined eDNA in cool temperate settings. By contrast, this study employs eDNA to survey the fish fauna in tropical Lake Bacalar (Mexico) with the additional goal of assessing the possible presence of invasive fishes, such as Amazon sailfin catfish. Sediment and water samples were collected from eight stations in Lake Bacalar on three occasions over a 4-month interval. Each sample was stored in the presence or absence of lysis buffer to compare eDNA recovery. Short fragments (184-187 bp) of the cytochrome *c* oxidase I (COI) gene were amplified using fusion primers and then sequenced on Ion Torrent PGM and S5 before their source species were determined using a custom reference sequence database constructed on BOLD. In total, eDNA sequences were recovered from 75 species of vertebrates including 47 fishes, 15 birds, 7 mammals, 5 reptiles, and 1 amphibian. Although all species are known from this region, 6 fish species represent new records for the study area, while 2 require verification. Sequences for five species (2 birds, 2 mammals, 1 reptile) were only detected from sediments, while sequences from 52 species were only recovered from water. Because DNA from the Amazon sailfin catfish was not detected, we used a mock eDNA experiment to confirm our methods were appropriate for its detection. We developed protocols that enabled the recovery of eDNA from tropical oligotrophic aquatic ecosystems, and confirmed their effectiveness in detecting diverse species of vertebrates including an invasive species of Amazon catfish.

## Introduction

Environmental DNA (eDNA) has gained popularity for biomonitoring, especially for the detection of invasive species and for baseline surveys of animal and plant communties [1,2]. This eDNA derives from cells shed into the environment as mucus, urine, feces, or gametes [2–7]. More than 120 articles have now considered eDNA, including a special issue on the topic [8]. The rapid adoption of eDNA-based bioassessments reflects two factors: improved access to the reference sequences [9,10] required to link the short reads from eDNA to their source species, and the availability of high-throughput sequencers that can generate large volumes of data at modest cost.

Conventional biomonitoring programs require extensive effort with sampling repeated over time and space [11]. Because these methods utilize nets and electrofishing, they are invasive and can damage habitats [3,12,13]. Furthermore, they rarely reveal all species in an environment, and each specimen that is collected must be identified to a species, requiring access to taxonomic experts [14,15]. Given these limitations, eDNA-based methods are an attractive approach for the quantification of biodiversity required for conservation and management [8,16].

Most prior eDNA studies have focused on the detection of invasive fish species [2,3,17] or rare/ endangered aquatic species [18,19] although they have also been used to examine the abundance and biomass of fish species [20,21]. Early results revealed complexities in data acquisition and interpretation that have provoked studies to improve sampling and DNA extraction while also minimizing methodological biases [16,22,23].

Varied protocols have been used to preserve eDNA, but they usually involve the immediate storage of samples at low temperature [21], precipitation [24], the addition of lysis buffers to filters [25], and the inclusion of surfactants [26]. While eDNA has traditionally been assumed to rapidly degrade, this is not always the case. For example, Deiner et al. [27] employed long-range PCR (>16 kb) to recover whole mitochondrial genomes from the six fish species in a mock community, and from 10 of 12 species represented in water samples from a stream. Prior studies have also demonstrated that eDNA metabarcoding is an effective alternative to traditional biomonitoring as it has revealed the species composition of fish communities in various ecosystems including a species-rich coastal sea in Japan [28] and a river in Indiana, USA [29]. In fact, in both cases, it revealed species that were overlooked by other sampling methods.

Although most past eDNA studies have examined temperate ecosystems, there is increasing interest in the use of this method in the tropics [30–32]. The present study sought to develop an eDNA protocol for biomonitoring the fish fauna of oligotrophic tropical Lake Bacalar and adjacent habitats, and to verify its capacity to detect an invasive fish, the Amazon sailfin catfish, *Pterygoplichthys* sp., which represents a serious threat in this region.

## Material and Methods

### Study site

Lake Bacalar, the largest freshwater habitat in the Yucatan Peninsula, is renowned for its striking blue color, the clarity of its water, and for the world’s largest occurrence of microbialites. With a length of more than 10 km [33–35], it occupies a larger fault basin [36] (50 km long, 2 to 3 km wide). With sediments derived from karst limestone, it represents the world’s largest freshwater lens system [37]. Lake Bacalar is not directly connected to the sea, but it shows a high rate of groundwater flow [38]. The northern part is connected to a complex system of lagoons including Laguna Chile Verde and Laguna Guerrero. The southern part of the lake has an indirect connection to the sea via the Hondo River that enters Chetumal Bay. There are four cenotes (sinkholes) in the lake whose depths range from 50-90 m.

Water temperatures in Lake Bacalar range from 25-28 °C, while the Secchi transparency averages 10.3 m. The lake is unstratified [33,39] with a water temperature of 27°C or higher throughout the year. Its surface waters are slightly alkaline (pH=7.8) with a conductivity of 1220 μS/cm [39] and HCO3 values higher than in marine environments [33].

All samples were taken from the littoral zone at a maximum depth of 0.5 m, except La Unión (2 m) and Xul-Ha (4 m). Eight sampling sites were examined along the main axis of Lake Bacalar and associated systems in the Hondo River (Fig 1). Samples were collected in December 2015, January 2016, and April 2016.

**Fig 1.**
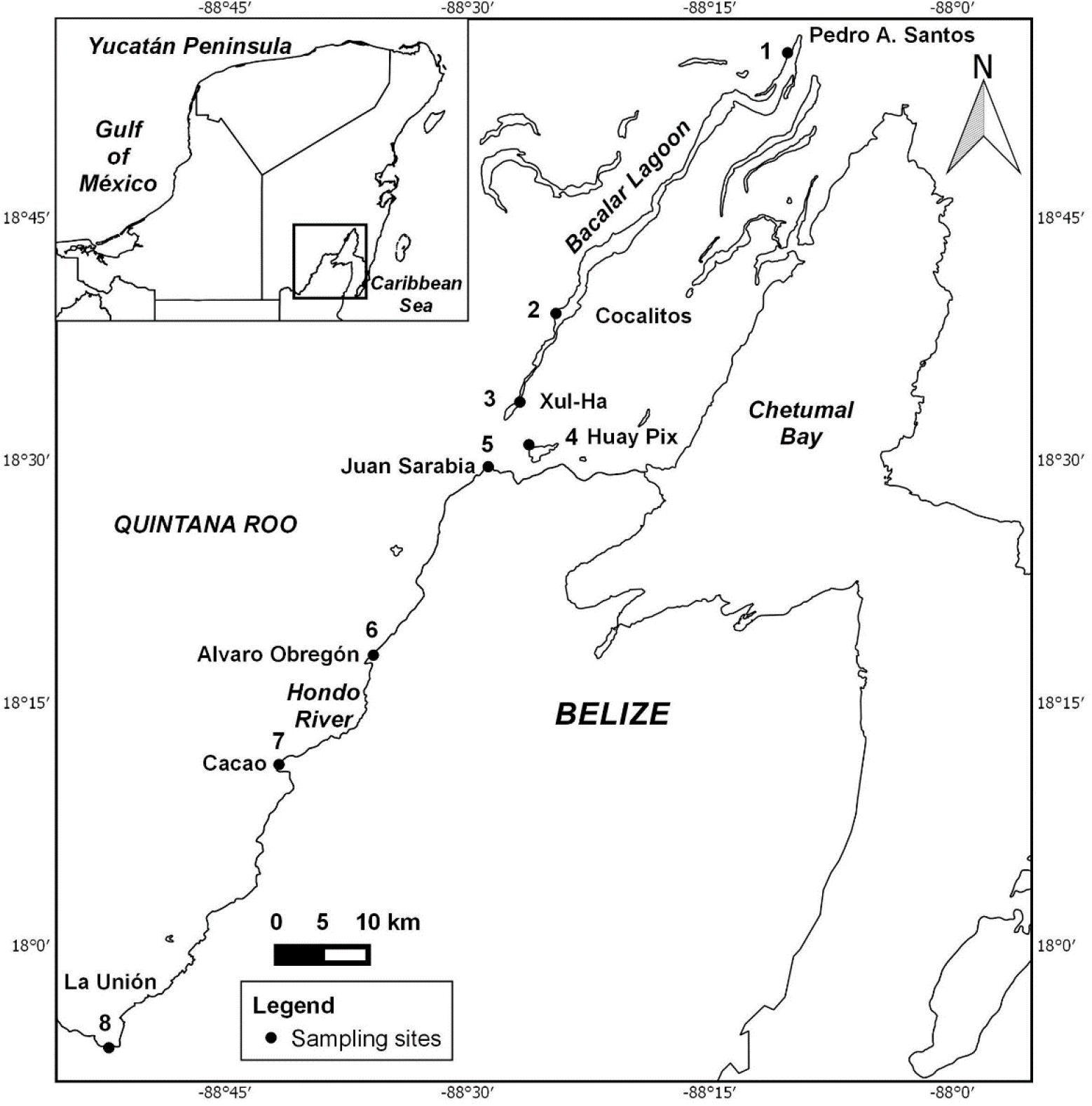
Sampling locations.

### Sampling

Eight localities (one with three points) (Fig 1) were sampled and most were examined on more than one occasion, producing a total of 14 sampling events. Three sites (Cocalitos, Huay Pix and Xul-Ha) were sampled in December 2015, four sites in January 2016 (Alvaro Obregón Viejo, Huay Pix, La Unión and Pedro A. Santos), and seven sites in April 2016 (Cacao, Cocalitos, Huay Pix, Juan Sarabia, La Unión, Pedro A. Santos and Xul-Ha).

All sampling bottles and equipment were handled with gloves to minimize contamination with human DNA. At each site, three replicate samples of water and sediment were taken. Each 1L water sample was placed in a sterilized CIVEQ^®^ bottle, while each sediment sample was placed in a separate (100 cm^3^) Ziplock bag. Each water and sediment sample was collected using a new turkey baster. Sediment samples were obtained from the upper 1 cm. All samples were immediately placed on ice, and then transported to the lab in Chetumal for processing.

### Processing water samples

Water samples were filtered within 7 hours of collection. Prior to processing, all lab surfaces and materials were sterilized with 10% bleach, followed by 70% ethanol; gloves were changed between each sample. Each sample was split into two 0.5 L subsamples that were filtered through separate 0.22 μm filters. One of these filters was stored with 1 ml of PW1 solution with grinding media while the other filter was placed with grinding media in a dry tube covered by aluminum foil. This approach resulted in six samples per locality for each collection event (Fig 2). All filters were stored at −18°C before being transported on ice from Chetumal to Guelph where DNA extraction was undertaken. The interval between filtration and DNA extraction was always less than 48 hours.

**Fig 2.**
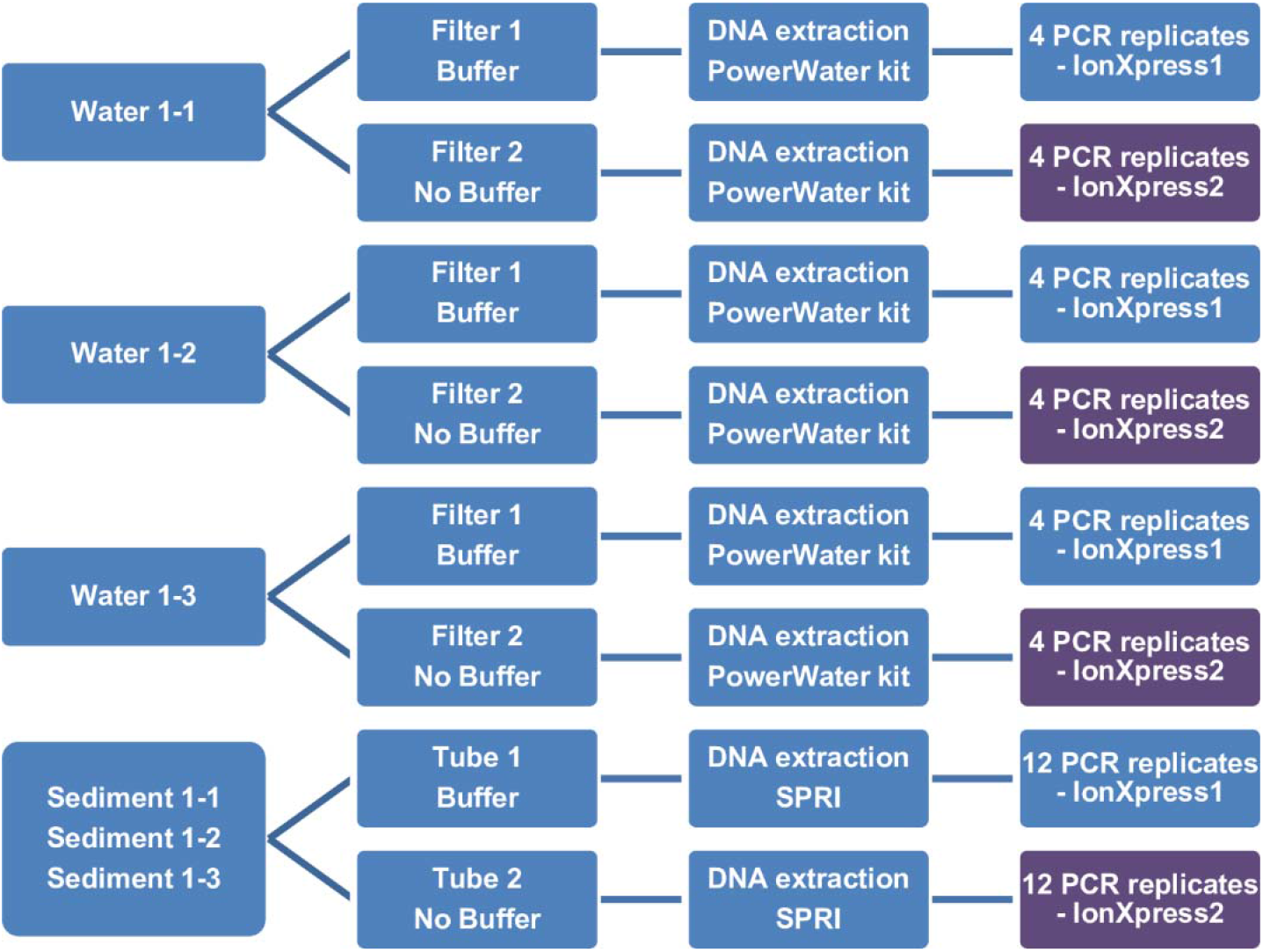
Experimental design for the water and sediment samples.

### Processing sediment samples

During the first collection trip, three replicate sediment samples (20 cm^2^ each) were collected at each site. Each sample was split into two 10 cm^3^ subsamples; 2 ml of 5M GuSCN buffer was either used as a preservation buffer or was added during lysis. For the second and third collection events, one pooled sample (60 cm^3^) was collected at each site and split into two replicates of 30 cm^3^; 6 ml of 5M GuSCN buffer [40] was used as a preservation buffer or was added during lysis step (Fig 2). Sediments were kept at -18°C before transportation on ice to Guelph.

### DNA extraction from water samples

Prior to DNA extraction, all lab surfaces and pipettors were sterilized using 10% bleach, followed by 70% ethanol. Pipettors were repeatedly cleaned with 10% bleach, followed by 70% ethanol during extraction and gloves were frequently changed. Centrifuge rotors, adapters, and tube racks were washed with diluted ELIMINase (Decon Labs) (1:10) and rinsed with deionized water.

DNA was extracted from the filters from the water samples using a PowerWater DNA extraction kit (MoBio) following a modified lysis protocol (tubes were incubated at 65°C prior to the grinding stage) with minor modifications to avoid cross-contamination between samples; all steps requiring 650 μl were reduced to 625 μl. Columns were incubated at 56°C for 15 min prior to DNA elution in 100 μl of PW6 buffer and quantified using Qubit 2.0 and HS dsDNA kit.

### DNA extraction from sediment samples

All lab surfaces and equipment were decontaminated as described above. The DNA extraction protocol employed ProK digestion in the presence of chaotropic salts followed by the use of magnetic beads to bind and then release DNA. Sera-Mag SpeedBead Carboxylate-Modified Magnetic Particles (Hydrophilic) from GE Healthcare were employed to bind/release DNA.

They were prepared as described in [41] in a polyethylene glycol/NaCl buffer (0.1% Sera Magnetic Particles (w/v) in 18% PEG-8000, 1M NaCl, 10 mM Tris-HCl, pH 8.0, 1 mM EDTA pH 8.0, 0.05% Tween 20). Extraction was initiated by adding a 1/5 volume of 5M GuSCN lysis buffer [40] to non-treated samples and a 1/10 volume of ProK (20 mg/ml). Each tube was vortexed and incubated for 2 hours at 56°C on an orbital shaker, then held for 30 min at 65°C before being centrifuged at 2000 g for 2 min. DNA was extracted from 500 μl of the resultant lysate after its transfer to a 2 ml Eppendorf LoBind tube. 500 μl of prepared magnetic beads was added to each lysate, and a pipettor was used to mix the beads and lysate before incubation at room temperature for 10 min. Tubes were then placed on a magnet (DynaMag) and incubated until the supernatant was clear. Beads were thoroughly washed once with 1 ml of PWB buffer [42] and twice with 1 ml of 70% ethanol. Magnetic beads were then dried at room temperature for 10-15 min before DNA was eluted into 100 μl of 10 mM Tris-HCl, pH 8.0 and quantified using Qubit 2.0 and HS dsDNA kit.

### PCR amplification

Prior to PCR amplification, each DNA extract from sediments was diluted 1:10 with molecular grade water while extracts from water were used without dilution.

In order to increase species recovery and overcome possible PCR bias introduced by UMI-labelled fusion primers [43] we followed a two-step PCR approach [43] with the first round employing conventional primers, where the diluted PCR product serves as a template for a second round of PCR with fusion primers containing sequencing adapters and UMI-tags.

A 184-187 bp segment of the barcode region of COI was amplified with two primer sets (AquaF2/C_FishR1, AquaF3/C_FishR1) known to be effective for vertebrates (Table 1). The PCR reactions employed the master mix described in [44] and Platinum Taq. The first round of PCR employed the following thermal regime: 94°C for 2 min, 40 cycles of 94°C for 40 s, annealing at 51°C for AquaF2 or 50°C for AquaF3 for 1 min, and 72°C for 1 min, with a final extension at 72°C for 5 min. Following the first round, PCR products were diluted 2-fold with molecular grade water, and 2 μl was transferred to a new tube for a second round of PCR to create amplicon libraries tagged with unique molecular identifiers (UMIs) with fusion primers containing Ion Xpress^™^ MID tags and Ion Adapters (Table 4). The same UMI-tag was applied to all three replicates for each treatment from a particular collection event (Fig 6). The PCR regime for the second round consisted of 94°C for 2 min, 20 cycles of 94°C for 40 s, annealing at 51°C for 1 min, and 72°C for 1 min, with a final extension at 72°C for 5 min. PCR products were visualized on an E-Gel96 (Invitrogen, Thermo Fisher Scientific).

**Table 1.**
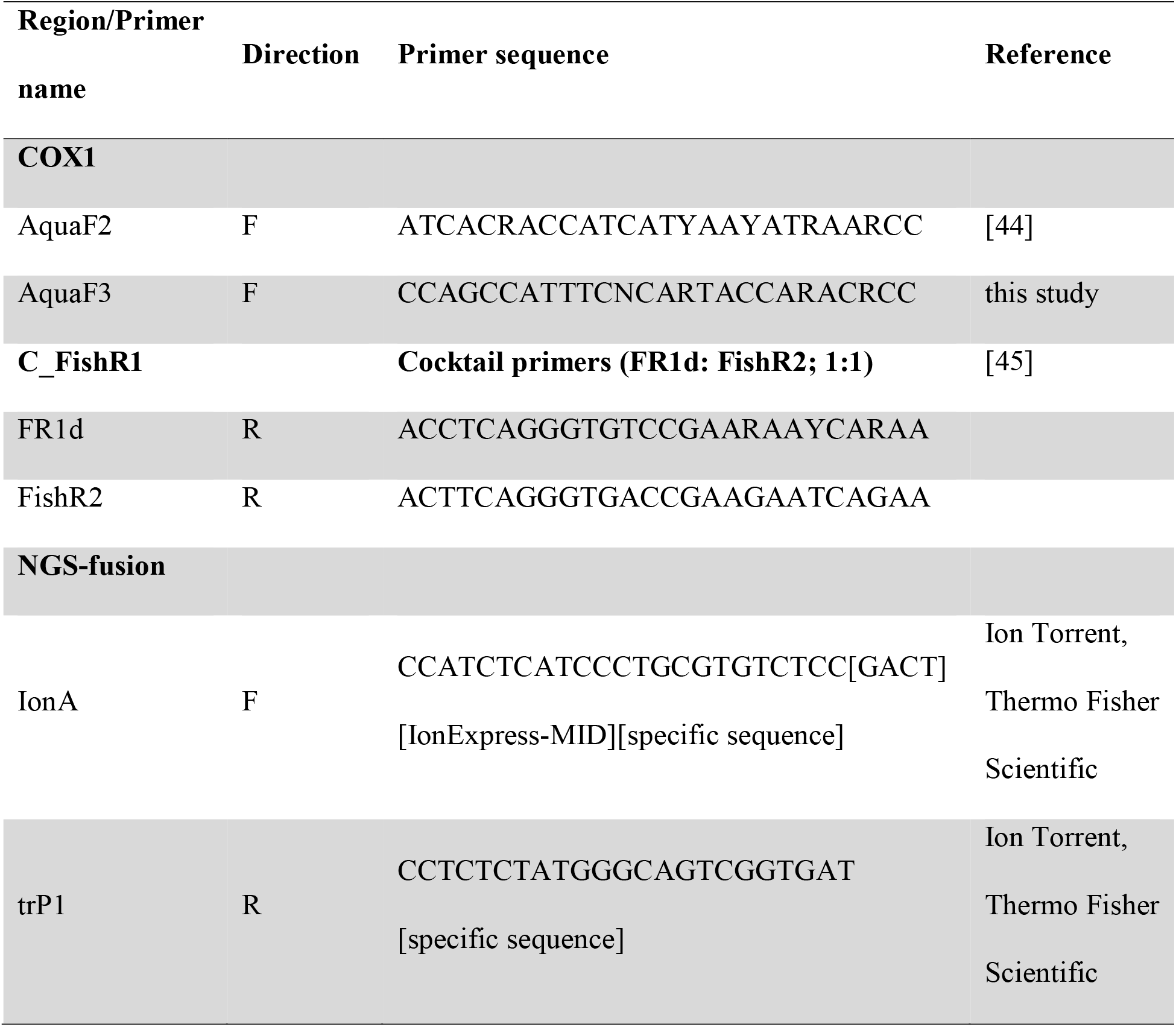
Primers used for eDNA amplification

Libraries for AquaF2/C_FishR1 and AquaF3/C_FishR1 were purified separately using magnetic beads prepared as described in [41], quantified using Qubit 2.0 and HS dsDNA kit, and normalized to 13 pM prior to sequencing. Most amplicon libraries were sequenced on an Ion PGM using the 400 Template and Sequencing kits following manufacturer’s instructions with either a 316 or 318 chip. Water samples from April 2016 were also analyzed on an Ion S5 using the Chef for templating and an Ion 520 and 530 kit with a 530 chip. Fig 7 summarizes the treatment for each water or sediment sample. HTS data (FASTQ files) are available in the European Nucleotide Archive (ENA) under submission number: PRJEB25950.

### Detection of an invasive fish

We conducted a mock eDNA experiment to ensure our methods could detect species of *Pterigoplichthys*. Two of these fishes were kept in a 10 L aquarium for 10 days at 25°C. One liter of water was collected and two 0.5 L subsamples were filtered through separate 0.22 μm filters; filters were stored with buffer and no buffer as described above.

### Bioinformatics workflow

We created a reference dataset (DS-EBACALAR) on BOLD with sequences from 3,534 Mexican fishes representing 576 BINs, including 219 records from Lake Bacalar and surrounding habitats (Hondo River, and adjacent lakes – Huay Pix, Xul-Ha). This reference barcode library included sequences for 53 of the 57 fish species (Table 2) known from these habitats (see Fig 1). As well, four other datasets with all public data on BOLD for Amphibia, Mammalia, Aves, and Reptilia were constructed. These five datasets were employed as a basis for assigning a source species to each OTU recovered from the present eDNA analyses.

**Table 2.**
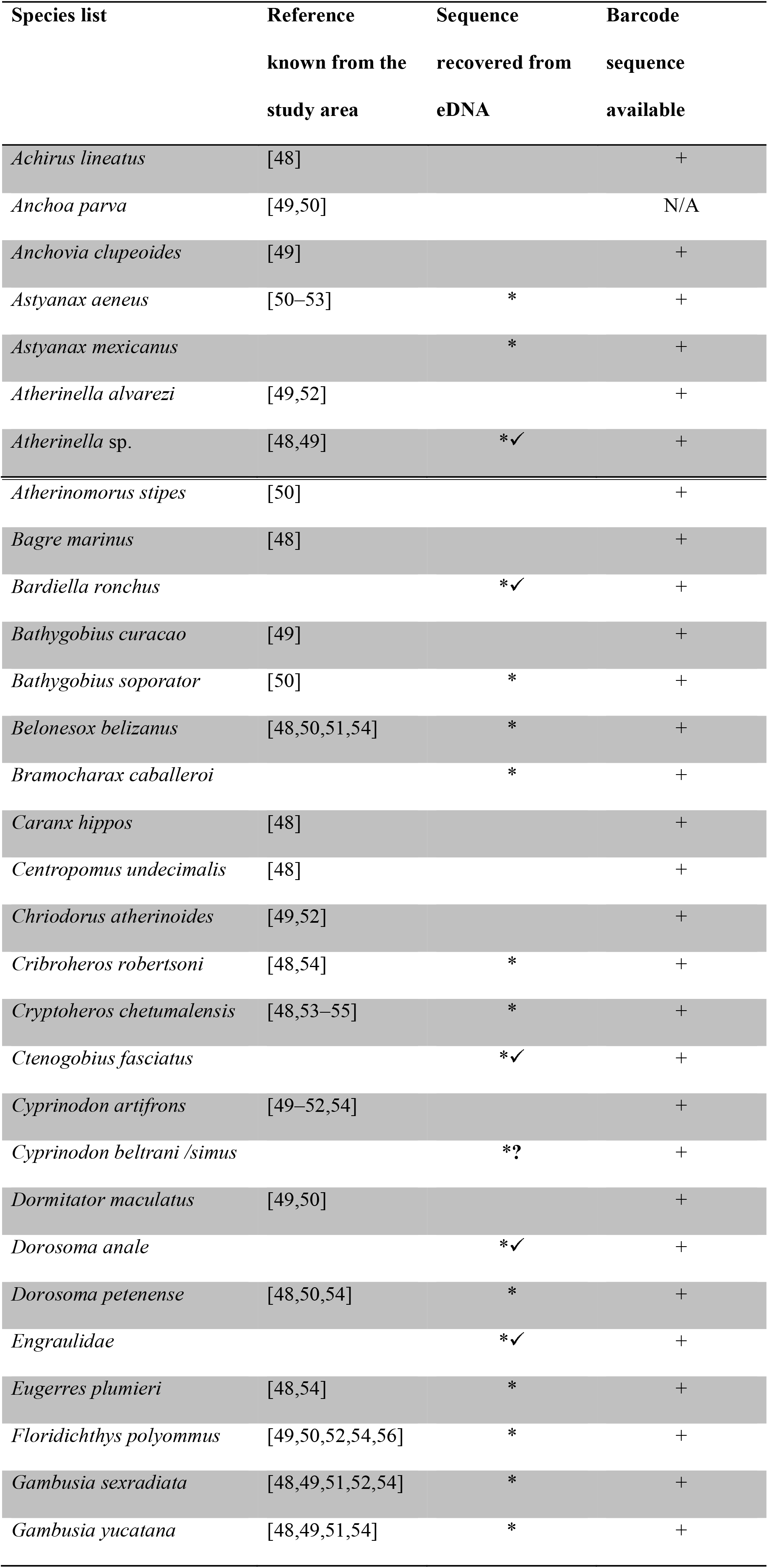
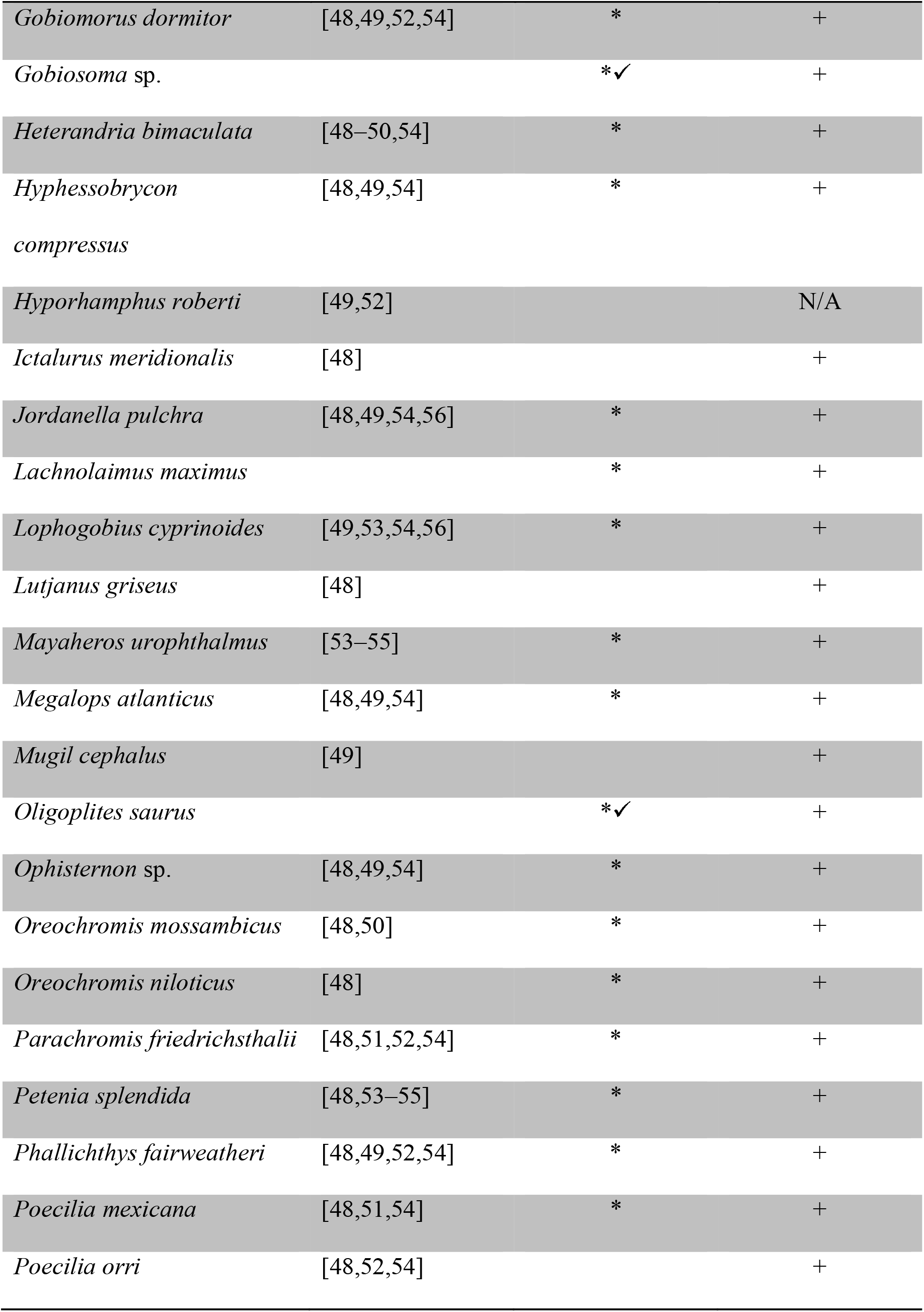
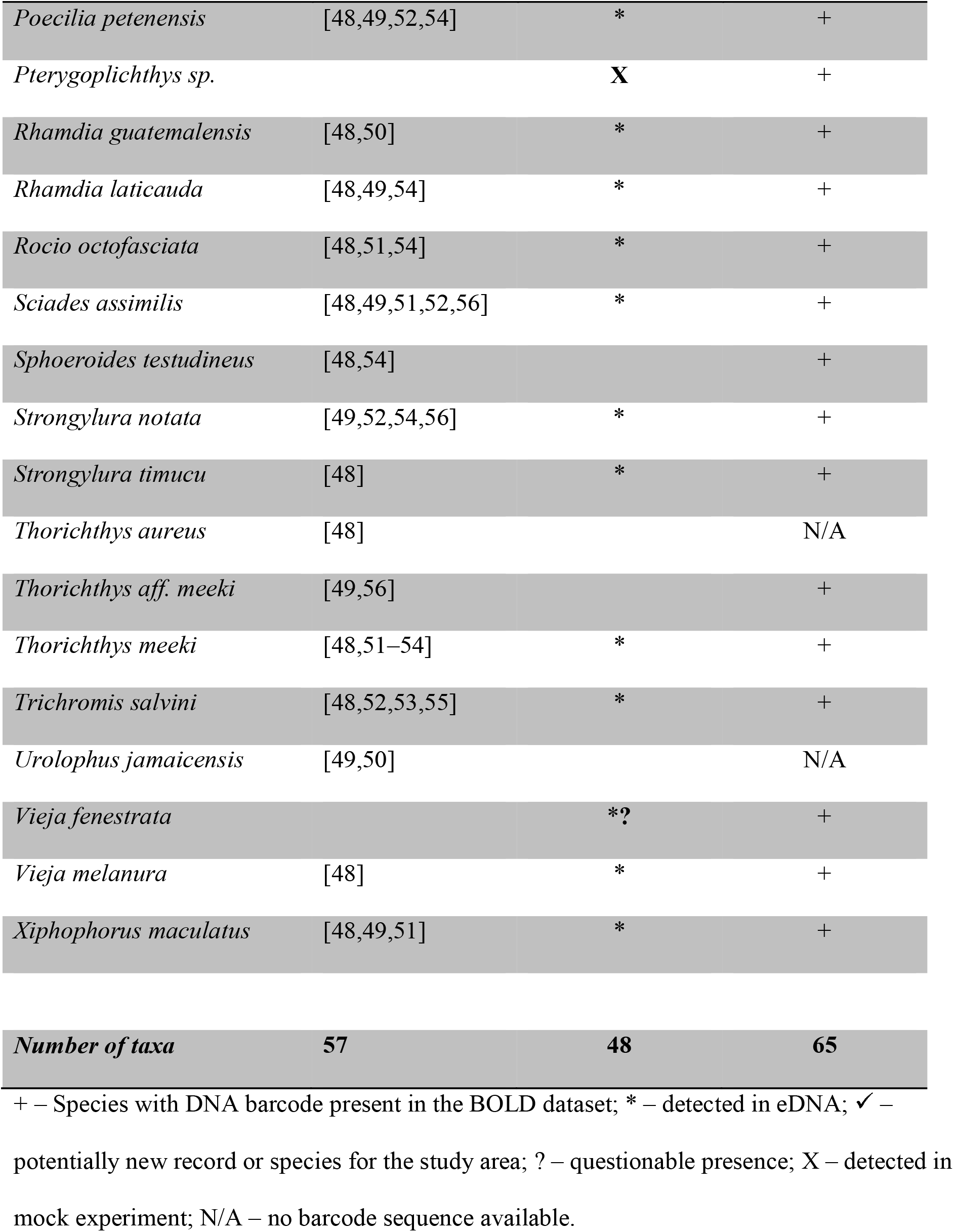
Fishes that were detected in eDNA, known from the literature and those with barcodes.

The following procedure was used to process the raw NGS reads: Cutadapt (v1.8.1) was used to trim primer sequences; Sickle (v1.33) was used for size filtering (sequences from 100-205 bp were retained), while Uclust (v1.2.22q) was employed to recognize OTUs based on the >98% identity and a minimum read depth of 2 thresholds. The Local Blast 2.2.29+ algorithm was then used to compare each OTU to the reference sequences in five datasets: DS-EBACALAR Bacalar Fish Dataset for eDNA Detection (3,534 sequences), and public BOLD data for Amphibia, Aves, Mammalia, and Reptilia represented by the following datasets: DS-EBACAMPH (11,018 sequences), DS-EBACAVES (28,914 sequences), DS-EBACMAMM (39,890 sequences), DS-EBACREPT (5,424 sequences). Raw Blast output results were parsed using a custom-built python script OTUBlastParser.py, and concatenated using ConcatenatorResults.py available at https://gitlab.com/kbessonov/OTUExtractorFromBLASToutput; processed results in tab-delimited format were imported to MS Excel, filtered by min score of 250, and 97-100% percent identity range. Filtered results werefurther processed and visualized in Tableau software 10.2. All human reads were excluded from the final results and species counts.

Species accumulation curves were calculated in EstimateS software available at http://purl.oclc.org/estimates [46], with the following settings: 1000 randomizations; do not extrapolate rarefaction curves; classic formula for Chao 1 and Chao 2. Results were exported to Microsoft Excel and visualized in Tableau software 10.4.

Sequences with 97-100% identity to fish species represented in the DS-EBACALAR dataset were mapped onto a neighbor-joining Taxon ID tree generated in BOLD for these taxa, and visualized in iTOL [47]. Results The eDNA analyses recovered sequences from 75 species of vertebrates including 47 fishes (Table 2), 15 birds, 7 mammals, 5 reptiles and 1 amphibian, most previously known from this region. Although substantially more species were recovered from the water than from the sediments (Table 3, Fig 3), five species were only recovered from sediments – racoon (*Procyon lotor*), pig (*Sus scrofa*), meso-american slider turtle (*Trachemys venusta*), brown jay (*Psilorhinus morio*), and great blue heron (*Ardea herodias*). However, all fish species detected in sediment samples were also recovered from the water samples (Table 3, S1 Table).

**Fig 3.**
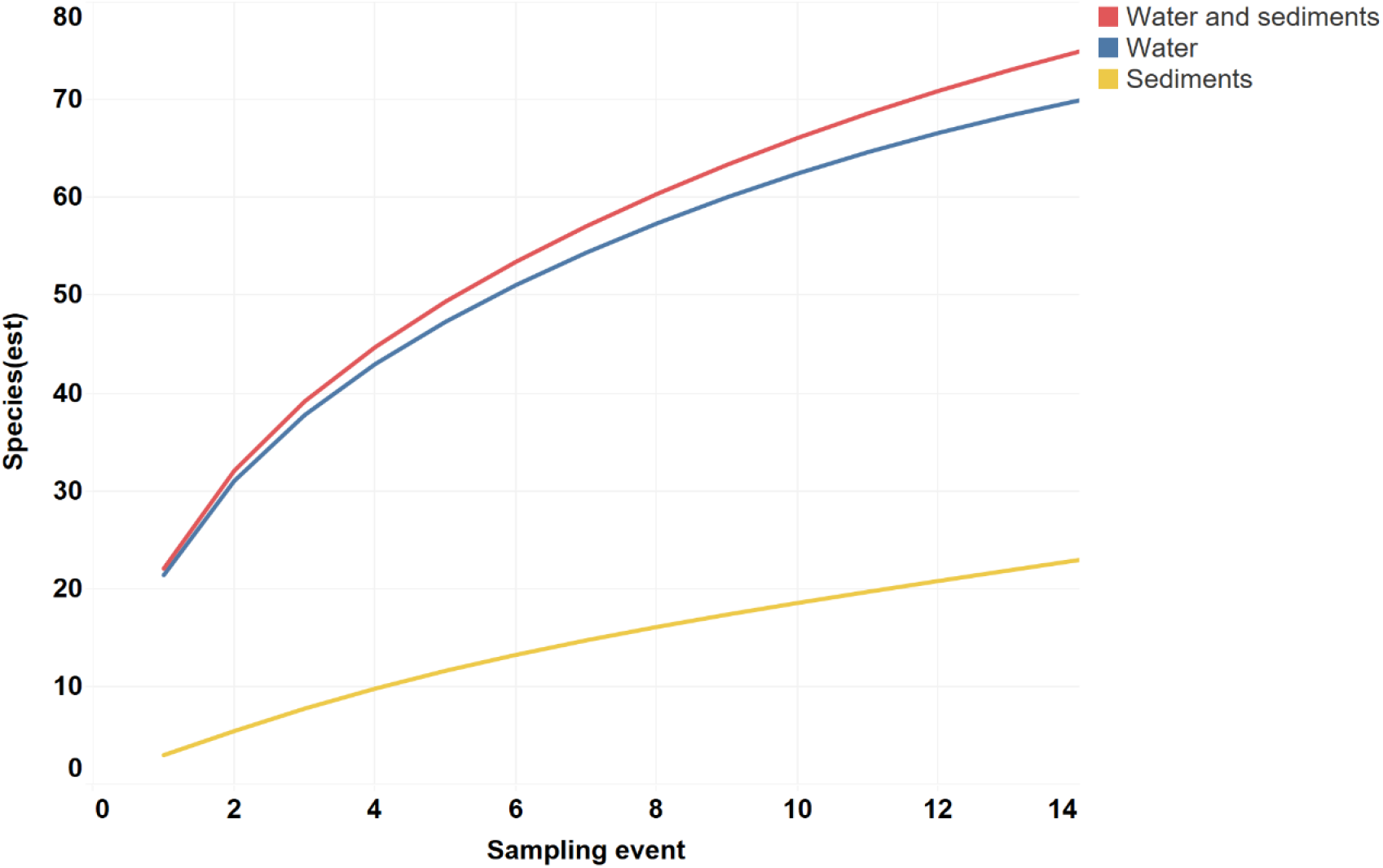
Number of species detected versus the number of samples analyzed for water, sediments, and combined. Species accumulation curves were calculated in EstimateS software with 1000 randomizations and classic formula for Chao 1 and Chao 2.

**Table 3.** Number of species recovered from the sediment and water samples.

|  | Total | Sediments only | Sediments + water | Water only |
| --- | --- | --- | --- | --- |
| **Actinopterygii** | 47 | 0 | 14 | 33 |
| **Amphibia** | 1 | 0 | 1 | 0 |
| **Reptilia** | 5 | 1 | 1 | 3 |
| **Mammalia** | 7 | 2 | 1 | 4 |
| **Aves** | 15 | 2 | 1 | 12 |
| **Total:** | **75** | **5** | **18** | **52** |

The three fish orders with highest number of detected species were Cichliformes (12 species), Cyprinodontiformes (11 species), and Gobiiformes (5 species) (S2 Table). Although *Pterygoplichthys* sp. was not recovered from water and sediment samples, it was recovered from the mock eDNA study (S1 Table and S2 Table).

In total, 272,610 reads obtained by the PGM from the water and sediment samples showed a close match to a vertebrate species in one of the five reference libraries. The three sites with the most fish species recovered were Alvaro Obregón Viejo, Cocalitos and Xul-Ha (Fig 4, Table 4), with 24-25 fish species per site. On average, the sediment and water samples from a particular site on a particular date generated eDNA sequences for 45% (mean =20.9; range= 15-25) of the 47 fish species detected in the survey (Table 4). Although nearly 2/3 of these reads derived from sediments (177,077 reads versus 95,533 from water), more species of vertebrates were recovered from the water samples (Fig 4). In fact, no species were recovered from the sediment samples taken at three sites (Cocalitos, Huay Pix, Xul-Ha) in December 2015 or from those preserved without buffer from Alvaro Obregón Viejo in January 2016. Although water samples revealed more species, there was evidence of temporal shifts in species recovery and DNA concentration with the lowest value in December 2015 (Fig 4, 5A). Best results were obtained with filtered water rather than sediments for both treated and untreated samples (Fig 3, 4, 5A).

**Table 4.** Number of vertebrate species detected at each of the eight sampling sites

| Sampling site No | Sampling site | Actinopterygii | Amphibia | Aves | Mammalia | Reptilia |
| --- | --- | --- | --- | --- | --- | --- |
| 1 | Pedro A. Santos / 18.920052°; -88.170100° | 18 |  | 2 | 1 |  |
| 2 | Cocalitos / 18.651469°; -88.408948° | 24 | 1 | 5 |  | 1 |
| 3 | Xul-Ha / 18.560084°; -88.446216° | 25 |  | 1 |  | 2 |
| 4a | Huay Pix / 18.514639°; -88.426750° | 19 |  | 3 |  |  |
| 4b | Huay Pix / 18.528561° -88.429242° | 15 |  | 1 |  |  |
| 4c | Huay Pix / 18.516144°; -88.436876° | 19 |  | 1 | 1 | 1 |
| 5 | Juan Sarabia/ 18.493471°; -88.478857° | 19 |  | 1 | 1 |  |
| 6 | Alvaro Obregón Viejo / 18.299259°; -88.597250° | 24 |  | 6 | 2 | 1 |
| 7 | Cacao / 18.186488°; -88.694636° | 23 |  | 1 | 2 |  |
| 8 | La Unión / 17.894676°; -88.870059° | 23 |  | 4 | 4 | 2 |

**Fig 4.**
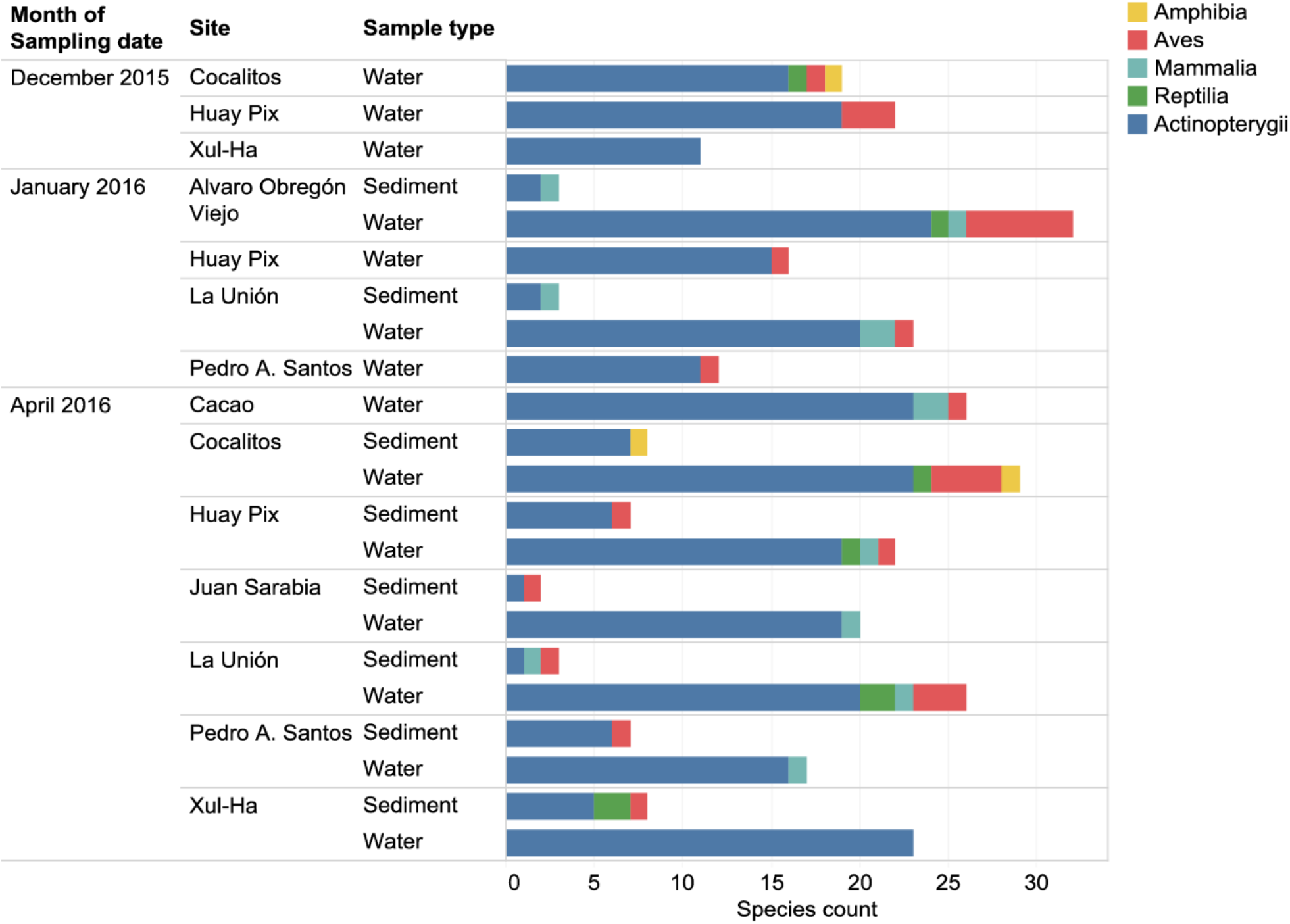
Number of vertebrate species recovered in eDNA from each sampling event at each collection site.

**Fig 5.**
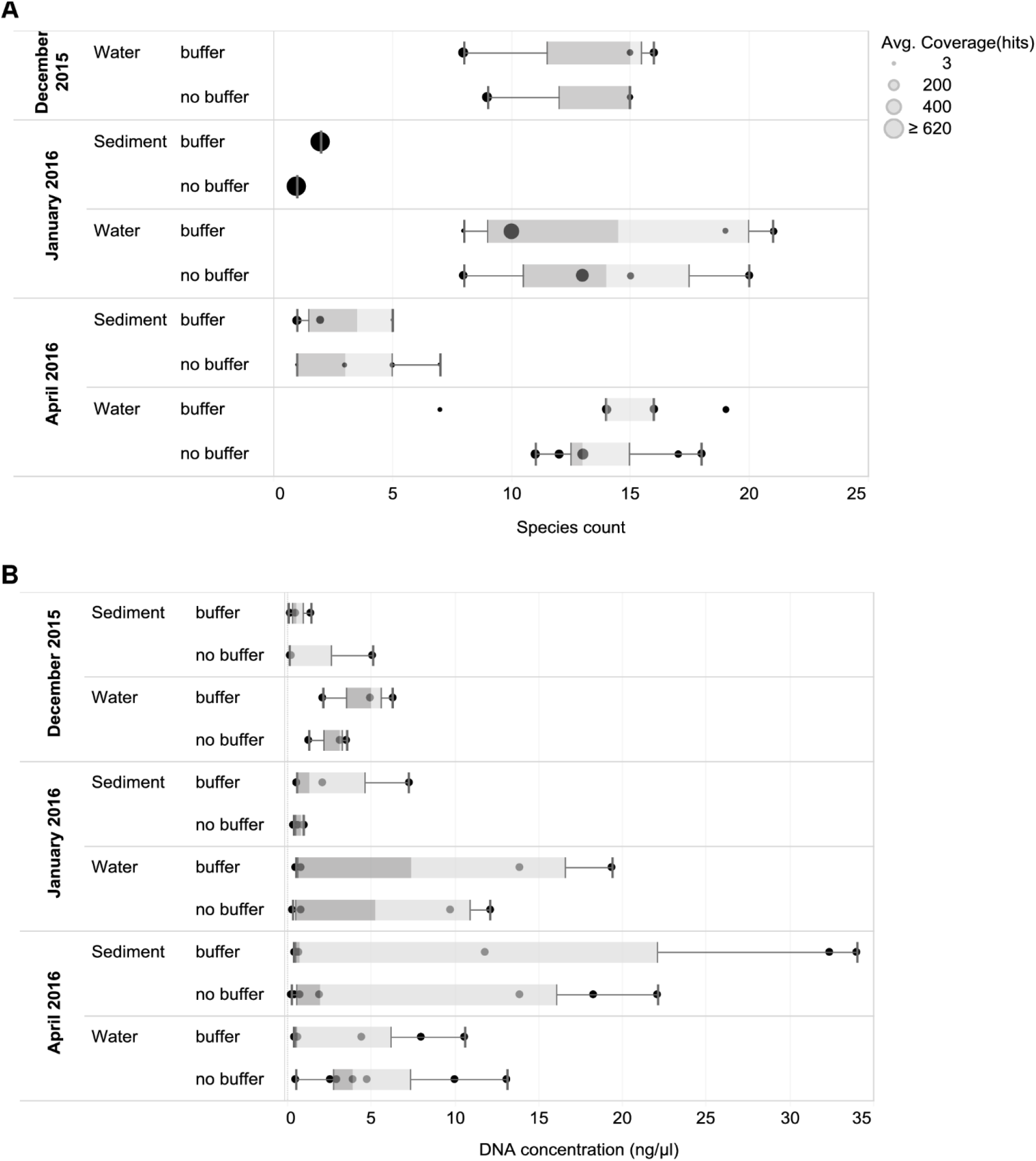
Species recovery and DNA concentration for each sampling event and treatment. A – number of vertebrate species recovered for each sampling event and treatment (only PGM data for fishes are shown, circle size indicates average read count per sampling event). B – DNA concentration of sediment and water samples (one replicate per sampling event) measured on Qubit. Whiskers correspond to data within 1.5x of interquartile range (IQR).

With two exceptions, negative PCR and DNA extraction controls did not produce any vertebrate reads aside from those deriving from humans (S2 Table). Two reads of *Trichromis salvini* were recovered from a negative control on the S5, and two reads of *Mayaheros urophthalmus* were obtained from a negative control on the PGM. Because both cases involved a species common in samples from the same run, they likely reflect tag-switching due to a misread UMI [57,58] or cross-contamination. Given their rarity, it is unlikely such analytical errors significantly impacted our overall conclusions.

### Comparison of results from two sequencing platforms

Two sequencing platforms (PGM, S5) were used to analyze the same set of water samples from April 2016 and from the mock experiment. The PGM generated 57,689 reads that matched vertebrates while the S5 generated 1,106,574 vertebrate reads. Reflecting its 20-fold higher sequence count, the S5 recovered more fish species than the PGM (41 vs. 34). However, large shifts in the relative number of read counts for the component species were also detected. For example, *Trichromis salvini* comprised 59.1% of the S5 reads, but just 5.9% of those from PGM (Fig 6).

**Fig 6.**
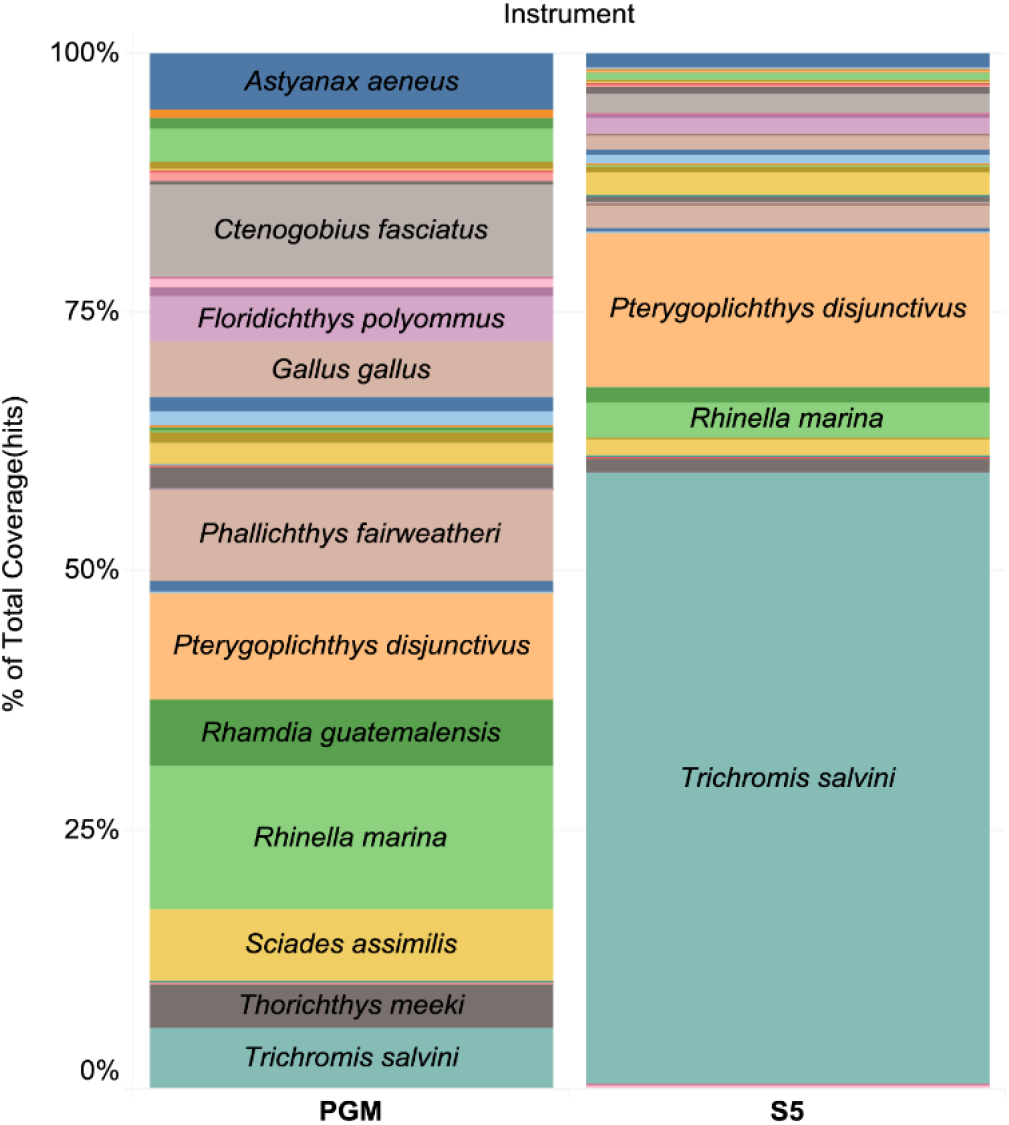
Percentage of total read coverage for each species recovered in water samples from April 2016 and mock experiment, analyzed with Ion Torrent PGM and S5 instruments. To verify the accuracy of the identifications assigned to the sequences recovered from eDNA, we generated a Neighbor-Joining tree in BOLD for detected fish species and mapped corresponding top-hit process IDs (Fig 7).

**Fig 7.**
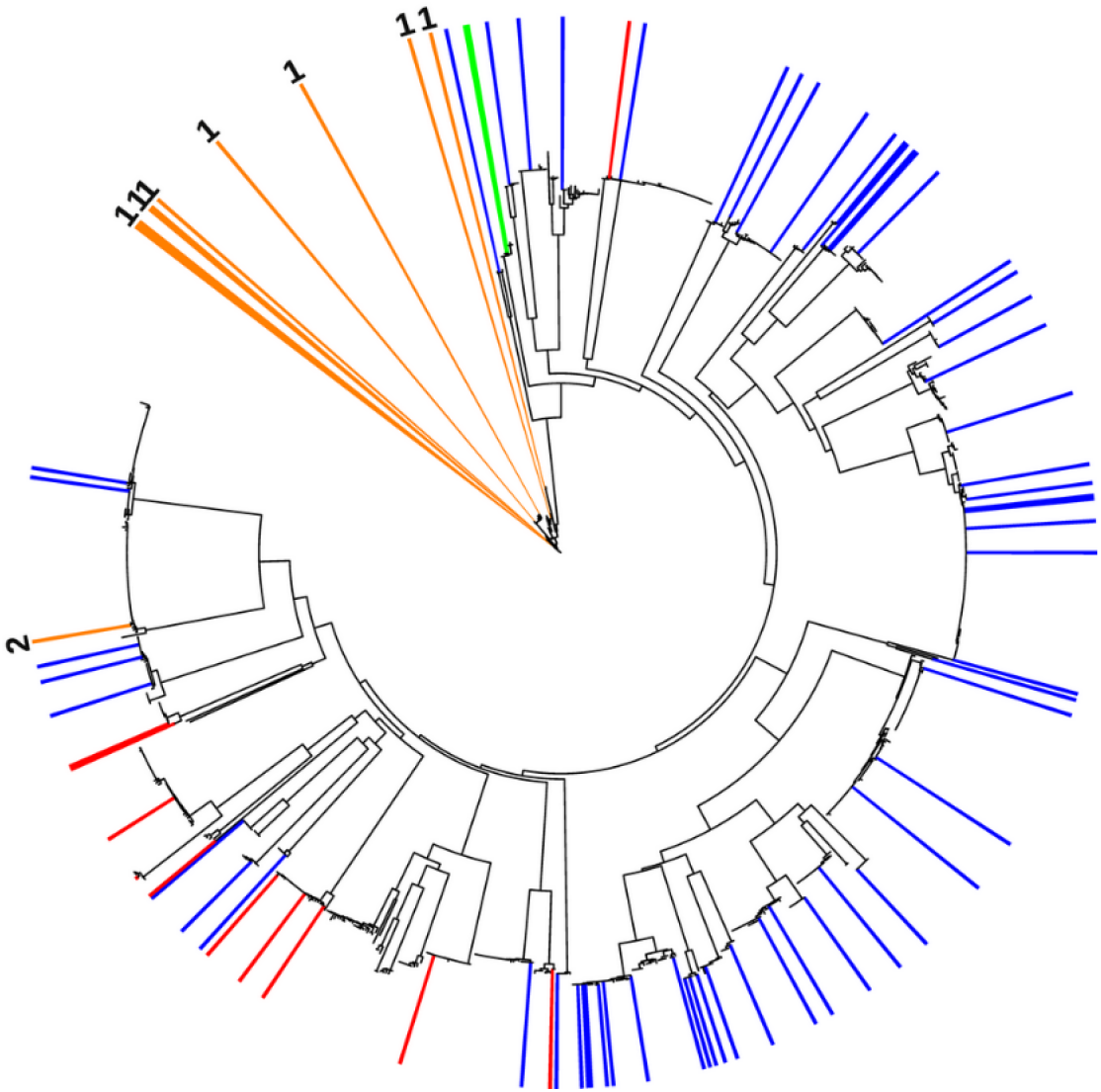
Top hit process IDs mapped on the Neighbor-Joining tree for fish taxa visualized in iTOL. Blue – species previously reported from the Lake Bacalar region; red – species new to Lake Bacalar); orange – species complexes lacking resolution with COI: 1 – *Bramocharax-Astyanax* complex, 2 – *Cyprinodon simus/beltrani;* green – *Pterygoplichthys* from mock eDNA experiment.

## Discussion

Although eDNA has often been thought to degrade rapidly, factors such as temperature, alkalinity, and trophic state [31,59] affect its stability. For example, cold temperatures, low UV-B levels, and high pH slow eDNA degradation [59], while acidity promotes it [59,60]. The overall probability of eDNA detection also depends on its production which may vary by species, by season, by density, and diet, and its loss from the study system via water discharge or diffusion [59].

Because techniques for eDNA analysis are still being optimized, some contradictory results have been reported. For example, a laboratory study showed that eDNA degradation increased with rising temperature, particularly in water samples from an oligotrophic lake [31]. By contrast, Robson et al [32] evaluated effects of high water temperature and fish density on the detection of invasive Mozambique tilapia in ponds via eDNA protocols and found that increased water temperatures did not affect degradation rates. However, they did detect increased rates of eDNA shedding at 35°C.

The present study represents the first time that the effectiveness of eDNA has been tested in an oligotrophic tropical lake, where high temperatures (>27°C year round) should speed DNA degradation. However, its water is slightly alkaline (pH 7.8) [39] which may aid eDNA preservation, perhaps explaining the better recovery of eDNA from water samples than sediments (Fig 2, 5A). The recovery of species from sediments were low (0-8 species per site) despite the higher DNA concentration in most samples treated with buffer (Fig 5B). The low recovery may reflect the presence of inhibitors or nuclease activity although best practices were followed to minimize DNA loss [61,62]. In particular, samples were transferred onto ice immediately following collection, and DNA was promptly extracted from the samples.

Most prior eDNA studies have examined vertebrates in temperate freshwater streams [63], lakes [64] or sea water [65,66]. However, Lopes et al. [30] recently demonstrated that eDNA surveys were effective in revealing anurans in streams in the Brazilian Atlantic forest, while Robson et al. [32] developed eDNA protocols that employ qPCR to detect invasive fish in tropical environments. Our results further validate the effectiveness of eDNA surveys in these settings.

There are three major approaches for of eDNA detection: the use of qPCR to detect one or a small number of target species [67,68], shotgun sequencing of aquatic metagenomes [69] and PCR-based metabarcoding [28,29,63–66]. Metabarcoding studies cannot deliver information on species composition without a reliable reference database, such as that available for Mexican freshwater fishes which includes records for 93% of the 70 species known from Lake Bacalar and associated wetlands [48,51–54,70]. The present study has affirmed the effectiveness of eDNA analysis as a tool for rapidly assessing the species composition of fish communities. The extraction of eDNA from just 42 liters of water collected on three dates at 14 sampling points revealed 41 of 57 of previously recorded species and added 6 potentially new species to the fauna of the Lake Bacalar region. The present results mirror those obtained in a study which used eDNA to examine the composition of fish communities a coastal sea [28]. The analysis of 94 water samples obtained by sampling 47 sites in six hours revealed 40 of the 80 species known from 14 years of underwater surveys, as well as 23 new species.

The present study recovered sequences records from six fish species new to the Lake Bacalar region (*Bardiella ronchus, Ctenogobius fasciatus, Gobiosoma* sp., *Dorosoma anale, Oligoplites saurus* and one Engraulidae). Two of the new species (*B. ronchus, O. saurus*) detected in Juan Sarabia and Alvaro Obregón Viejo are known from the adjacent Chetumal Bay, but have not previously been reported from inland waters (Valdez-Moreno, pers. obs.). The new gobiid, *C. fasciatus*, was detected in four localities (Xul-Ha, Pedro A. Santos, Cocalitos, Huay Pix), but was overlooked in prior field surveys because of its morphological similarity to other gobids in Lake Bacalar, especially *Lophogobius cyprinoides*. The other two new records require confirmation – the presence of a *Gobiosoma* in Huay Pix (51 reads) and a member of the family Engraulidae in Pedro A. Santos (>7000 reads).

A few sequences (13) of *Lachnolaimus maximus*, a commercially fished marine species, were recovered from Huay Pix, but the presence of its DNA was almost certainly mediated by human activity as fish were prepared for consumption.

As reported previously, species in two characid genera (*Astyanax* and *Bramocharax*) are difficult to discriminate using DNA barcodes as they show low divergences between species and genera [51,71]. Moreover, in a study utilizing three mitochondrial genes (Cyt b, 16S, COI) and a single nuclear gene (RAG1), *Bramocharax* was found to be polyphyletic, with species in this genus being sisters to different clades of *Astyanax*, making *Astyanax* paraphyletic [72]. In fact, all *Bramocharax* species grouped with sympatric *Astyanax* lineages (or even with allopatric *Astyanax* populations), with less than 1% divergence. In our dataset, *Astyanax aeneus, Astyanax mexicanus*, and *Bramocharax caballeroi* also formed an intermixed cluster in the Taxon ID tree generated (see the dataset DS-EBACALAR in www.boldsystems.org). Among the species detected with eDNA, *Astyanax mexicanus* was the least certain as it was only represented by 7 reads from the S5 with an average identity of 0.97 (S2 Table), which may be low quality reads.

Although *Dorosoma anale* was only represented by 13 reads from water at La Unión station, this species is known to occur at sites in northern Belize close to the Hondo River (Valdez-Moreno pers. obs.).

The presence of two other species (*Cyprinodon beltrani* (two PGM reads) and *Vieja fenestrata* (five S5 reads) was less certain. *C. beltrani* is native to Chichancanab lagoon, and cannot be distinguished from *C. simus* with DNA barcodes (see DS-EBACALAR in www.boldsystems.org), while *V. fenestrata* is native to the Papaloapan River [52] so its presence in the study area is unlikely.

Our results also revealed good eDNA recovery for other vertebrates, including rare species, such as *Tamandua mexicana*, which was represented in both PGM and S5 sequences from a river sample (Juan Sarabia). Another interesting record involved *Oreothlypis peregrina*, a migratory warbler that is known to overwinter in the Yucatan Peninsula, but that is very morphologically similar to other species.

Biodiversity in freshwater ecosystems in undergoing losses as a result of landscape transformation, pollution, and biological invasions. In fact intentional or accidental introduction of invasive alien species is the third leading cause of global biodiversity loss [73]. In 2008 it was estimated that nearly 40% of the freshwater fish species in North America, including Mexico, were threatened by invasive species [74]. The Lake Bacalar water basin has been recognized as one of the hydrological basins with high priority for conservation by the Mexican National Commission for the Use and Knowledge of the Biodiversity (CONABIO) [75]. Fortunately, most wetlands and lakes in this region are relatively pristine, excepting the Hondo Riveron the border between Mexico and Belize, which has been heavily impacted by the discharge of organic waste and pesticides, by vegetation clearing, and by the introduction of invasive such as tilapia [76], and the Amazon sailfin catfish (*Pterygoplichthys pardalis*), which was recently detected [77,78] near La Unión [79]. The impact of this declining water quality and rising incidence of invasive species on the native fish fauna needs to be carefully monitored and eDNA-based studies could provide a cost-effective way to meet this need

## Conclusions

We developed field sampling protocols and a HTS pipeline which enabled the efficient recovery of eDNA from several tropical aquatic ecosystems. Water samples consistently revealed more vertebrate species than sediment samples although about 10% of the species were only recovered from sediments. eDNA sequences were recovered from 75 species of vertebrates including 47 species of fishes, with six new records for the Lake Bacalar region and two other species whose detection is likely to be due to human activity. Sequences were also detected from another 28 vertebrate species including 15 birds, 7 mammals, 5 reptiles, and 1 amphibian, all species known from the watershed.

This study indicates that eDNA can aid conservation and monitoring programs in tropical areas by improving our capacity to map occurrence records for resident and invasive species. There remains a need to convince both regulatory agencies and the public that this approach can provide the detailed information on species composition needed to underpin conservation policy for tropical aquatic ecosystems.

## Acknowledgments

We thank Jose Angel Cohuo, Miguel Valadez, and Rosaura Castro from the Instituto Tecnológico de Chetumal and Fernando Cortés Carrasco from El Colegio de la Frontera Sur for help with collecting and processing samples. DNA extraction and NGS were carried out at the Centre for Biodiversity Genomics (CBG), University of Guelph through funding to PDNH from Ann McCain Evans and Chris Evans. We thank Thomas Braukmann for designing the bioinformatics pipeline for processing the raw data. This paper represents a contribution from the Chetumal node of the Mexican Barcode of Life (MEXBOL) network. MVM and MEG thank CONACYT for support through the Sabbatical Stays Program (Grants 261790 and 262267).

## Author contributions

Conceived and designed the experiments: MVM, MEG.

Coordinated the collection of material and performed field experiments: MVM, MEG.

Performed the lab experiments: NVI, SLP.

Analyzed the data: NVI, MVM, MEG, SLP.

Designed illustrations: NVI, MVM.

Designed *OTUBlastParser.py* and *ConcatenatorResults.py* Python scripts: KB.

Wrote the paper: MVM, NVI, MEG, SLP, PDNH.

All authors have read and approved the final manuscript.

## Data accessibility for BOLD datasets

dx.doi.org/10.5883/DS-EBACALAR

dx.doi.org/10.5883/DS-EBACAMPH

dx.doi.org/10.5883/DS-EBACAVES

dx.doi.org/10.5883/DS-EBACREPT

dx.doi.org/10.5883/DS-EBACMAMM

## Supporting Information

**S1 Table. Supplementary Table 1.** Summary of coverage for vertebrate reads for all collection events and for positive controls.

**S2 Table. Supplementary Table 2.** Taxonomy, top hit Process IDs, average overlap, average score, coverage, and average identity for vertebrate species detected by eDNA in environmental samples, positive and negative controls.

